# InDeep : 3D fully convolutional neural networks to assist in silico drug design on protein-protein interactions

**DOI:** 10.1101/2021.07.28.453974

**Authors:** Vincent Mallet, Luis Checa Ruano, Alexandra Moine Franel, Michael Nilges, Karen Druart, Guillaume Bouvier, Olivier Sperandio

## Abstract

**Motivation:** Protein-protein interactions (PPIs) are key elements in numerous biological pathways and the subject of a growing number of drug discovery projects including against infectious diseases. Designing drugs on PPI targets remains a difficult task and requires extensive efforts to qualify a given interaction as an eligible target. To this end, besides the evident need to determine the role of PPIs in disease-associated pathways and their experimental characterization as therapeutics targets, prediction of their capacity to be bound by other protein partners or modulated by future drugs is of primary importance.

**Results:** We present `InDeep`, a tool for predicting functional binding sites within proteins that could either host protein epitopes or future drugs. Leveraging deep learning on a curated data set of PPIs, this tool can proceed to enhanced functional binding site predictions either on experimental structures or along molecular dynamics trajectories. The benchmark of `InDeep` demonstrates that our tool outperforms state of the art ligandable binding sites predictors when assessing PPI targets but also conventional targets. This offers new opportunities to assist drug design projects on PPIs by identifying pertinent binding pockets at or in the vicinity of PPI interfaces.

**Availability:** The tool is available on GitHub^3^ along with a PyMol plugin for visualization. Predictions of InDeep can be consulted at iPPI-DB^4^

## 1 Introduction

### Protein-protein interactions as therapeutic targets

Protein-protein interactions are central elements in numerous biological pathways. They represent increasing interests as therapeutic targets with a growing number of published studies describing the successful modulation of PPIs using small molecules [1]. Yet, identifying chemical probes or drugs on PPIs remains a difficult task. As opposed to more conventional drug discovery targets, such as G-protein coupled receptors (GPCRs) or enzymes and more recently protein kinases, PPIs have not evolved to bind small molecules. Therefore, the proof of their ligandability has to be made on a case by case scenario [2]. Indeed, the design of small molecules binding orthosterically at the interface to prevent protein interactions is not achievable for all PPIs [3]. The design of small molecules as allosteric or interfacial modulators, respectively binding distantly from the interface or in its vicinity, is sometimes a more appropriate strategy. Conversely, the design of therapeutic peptides or even antibodies might be inevitable solutions if the interface of the studied PPI is incompatible with the design of a small molecule. Given their number and heterogeneity of structures, it is therefore of primary importance to have powerful tools to efficiently evaluate the feasibility of considering PPIs as targets in complement of the unavoidable biological evaluations.

In the situation of designing orthosteric inhibitors of PPIs using small molecules (**iPPIs**), strategies like epitope mimetics can be envisaged [4]. This was successfully made against the B-cell lymphoma-2 (Bcl-2) family to combat chronic lymphocytic leukaemia[5]. This led to the development of Venetoclax which was approved by the FDA in 2016 as the first orthosteric PPI drug. The design of iPPIs as orthosteric epitope mimetics, allosteric or interfacial modulators implies to evaluate two complementary features within the interface: 1) the knowledge of an epitope binding at the interface and the presence of hotspot residues that carry out most of the binding energy of interaction [6], and 2) the existence of a ligandable binding site around these hotspots that could host a small molecule.

#### Predicting and profiling epitope binding sites

Alanine scanning is experimentally sound to evaluate the presence of hotspots but supposes the design of numerous point mutations, a process that is tedious and costly in particular when no partner is known yet. Several *in silico* tools can predict hot spot residues [7, 8] but they necessitate the structure of a complex and the fore knowledge of an identified protein partner. To predict protein interactions, some tools directly use the sequence information [9] or evolutionary data [10]. But leveraging the structure of the protein has shown to drastically increase performance of prediction of interface regions. Moreover, structural motifs and local arrangements of atoms can be highly conserved even across different secondary structures and different global protein folding. These local motifs are hypothesized to be the key element of partner binding. This has motivated using convolutional strategies to encode such local information about binding sites. Some of these methods focus on predicting the interaction patch on the protein : i.e. determine which residues are involved in an interaction [11, 12, 13, 14]. Some methods also take as input the partner to predict the interaction patch[12], deemed as partner-specific predictions. In that case, the prediction can be more fine-grained and also provide contact prediction : which residue interacts with which other [15, 16, 17, 18]. All of these tools annotate the sequence by predicting residue-level information. However, knowing not only which residues are involved in the binding (sequence or surface derived info) but which types of partner residues and where they bind in the vicinity of the protein surface is highly desirable to understand the mechanisms of epitope binding or the design of future drugs mimicking these epitopes.

#### Predicting ligandable binding sites within PPIs

A plethora of tools is now available to predict binding sites and binding site ligandability. One can cite historical and efficient geometric based methods such as Fpocket [19], VolSite [20] and mkgridXf[21], fragment-based methods like FTMap [22] and more recent and powerful methods using deep learning such as DeepSite [23], P2rank[24], Kalasanty[25], DeepSurf[26], or OctSurf[27], with Kalasanty being a reference in the field. These methods have demonstrated their predictive capacity to identify ligandable binding sites on conventional drug targets. Yet, none of the methods cited above is specific to the ligandability of PPIs, although FTMap was the first method to rise the question of PPI ligandability without really providing a ligandability score. Nonetheless, the specificity of PPIs with regard to ligandability is an important aspect to keep in mind when designing drugs. Indeed, interfaces of PPIs are historically described as rather flat, large and devoid from deep binding pockets [28, 29, 30]. Yet, there are numerous examples of co-crystallized orthosteric iPPIs (see https://ippidb.pasteur.fr/targetcentric/). It is therefore most legitimate to anticipate a specific expression of ligandability in the case PPIs. Using machine learning and PPI-specific data sets, one can exploit local information about PPI and PPI/drug interaction using 3D fully convolutional approaches.

#### Capturing holo-likeness along MD trajectories

PPIs are known to undertake important conformational changes depending on their binding state: apo, holo with ligand or a protein partner. These conformational changes affect the shape and binding capacity of interface binding pockets [31]. It is therefore of primary importance to take these conformational changes into account when profiling epitope and ligandable binding sites [22] as those will condition binding to partners. This represents a major challenge when attempting to identify chemical probes, using *in silico* methods such as virtual screening [32] or designing epitope mimetics, in the absence of the partner bound. Indeed, it is for example key for such methods to sample and identify so-called holo-like conformations prior to virtual screening in the context of ensemble docking[33, 34]. Methods have already been developed to monitor ligandability along molecular dynamics trajectories using geometric [35], or Deep Learning approaches [36, 37], although none of these are specific to PPIs. Moreover, no method is available to monitor interactability patches and epitope binding sites along molecular dynamics trajectories.

#### Contribution

Our work builds upon our last release of iPPI-DB [1] and of its new target-centric mode^5^, and aims to facilitate the identification of iPPIs. Our tool `InDeep` has capitalized on iPPI-DB structural data to train predictive models relying on 3D fully convolutional (3D-FCN) neural networks. It is a unified multi-tasking prediction tool that uses the 3D structure of proteins to predict ligandable binding for iPPIs and so-called interactability patches for epitope binding. We show that `InDeep` outperforms the state of the art of binding pockets detection methods and that our tool is especially efficient to detect iPPI and epitope binding sites. While remaining competitive on annotating the protein sequence with interactability, it also predict the spatial location of its putative partner. Our tool also enables tracking of these druggability and interactability scores for a given detected pocket along molecular dynamics trajectories. It is integrated in a PyMol [38] plugin (see Section D) for easy visualization of the predictions, making it a real toolbox for iPPI drug design. It is freely available at https://gitlab.pasteur.fr/InDeep/InDeep. Finally, the results of `InDeep` predictions can be consulted on the iPPI-DB website for every heterodimers and iPPI-bound proteins in the database at https://ippidb.pasteur.fr/targetcentric/.

## 2 Methods

### 2.1 Data curation and splitting

For training and assessing the 3D-FCN models, we have used the dataset that is presently accessible on the website of our database iPPI-DB (https://ippidb.pasteur.fr/targetcentric/). The dataset relies on two subsets: one contains Hetero-Dimeric complexes (`HD` interactions) and the other iPPI-bound protein complexes (`PL` interactions) where ligands bind one of the two partners at the interface within a `HD` complex (orthosteric inhibitors). These data were retrieved from the PDBe database [39] using a filtering procedure and different quality control checks. The procedure operates directly on a JSON file containing all available information within PDBe to select PDB structures accordingly. Ligands were selected as to be not more than 6 Å away from the interface and contain at least five heavy atoms. Heavy atoms within compounds must be one the following (C, N, O, S, P, I, B, Br, Cl, F). For Xray structures and CryoEM structures, resolutions should be below 3.5 Å. For Xray structures R_Free_-R_Factor_ were below 0.07. For CryoEM structures, FSC was below 0.143. Once the selected PDB structures were downloaded, all `PL` structures were superimposed onto the corresponding `HD` complexes (sharing an identical UNIPROT ID) in order to identify orthosteric compounds competing for the same interface. Finally, we removed structures with alternative locations for residues at the interface within `HD` or `PL` complexes. Thus, we built a dataset of 2290 `PL` complexes and 6736 `HD` complexes for the training and the assessment of the 3D-FCN models.

We have then split this data based on CATH[40] folds to avoid any structural overlap between our train, validation and test data sets. To do so, we have used the pre-computed annotations available for each protein domains and annotated our interaction domains with those CATH classifications. We have then put only specific CATH in the train, validation and train split as is listed in Supplemental (section A).

### 2.2 Data representation

Equipped with these sets of co-crystallized proteins with either a protein or ligand partner, we wish to represent it in a way a neural networks can learn on. We follow the volumetric CNN framework used, for instance, in [23, 25]. This framework consists in treating the interaction sites as 3D images. First we introduce functional atom types : α-carbon (Cα), donor and receptor of h-bonds and positively and negatively charged atoms and hydrophobic/aromatic atoms. At each voxel, we associate a vector that represents the probability of presence of these functional atom types. This vector is computed by considering the input protein as a sum of Gaussian densities centered around each atom in the dimension corresponding to each atom type. We then perform an interpolation of this Gaussian sum on a regular grid, which yields a vector at each grid point. To encode the protein partner, we start with the same encoding and add a ’void’ channel so that on each grid point the encoding vector sum to one and represents a probability distribution. To represent iPPIs, we use only one channel for all atom types. This encoding is dependent on the arbitrary basis in which the protein is represented. To limit this arbitrary choice, we applied rotational data augmentation during the training of the network.

### 2.3 Model architecture and learning

We now want to build a model that takes a protein structure as input and predicts iPPI ligandability (`PL` interaction) as well as protein interactability (`HD` interaction). These two machine learning tasks can benefit from the concept of multitasking, a well-documented phenomenon that means that one hybrid machine learning model that solves two tasks usually performs better than two separate ones [41, 42, 43]. Indeed, in the multitasking setting, each task benefits from the representations learnt using the other task’s supervision.

We use a U-Net [44] architecture for our prediction, with blocks containing 3D convolutions, Batch Normalization [45] and ELU activations [46]. After several shared layers, the network is split into a `PL` and an `HD` branch. These branches are two independent sequences of layers with a sigmoid and softmax activations for `PL` and `HD` respectively. A visual representation of this branching scheme is available in Figure 1 and a detailed description of the model is available in the Supplemental (section B).

**Figure 1:**
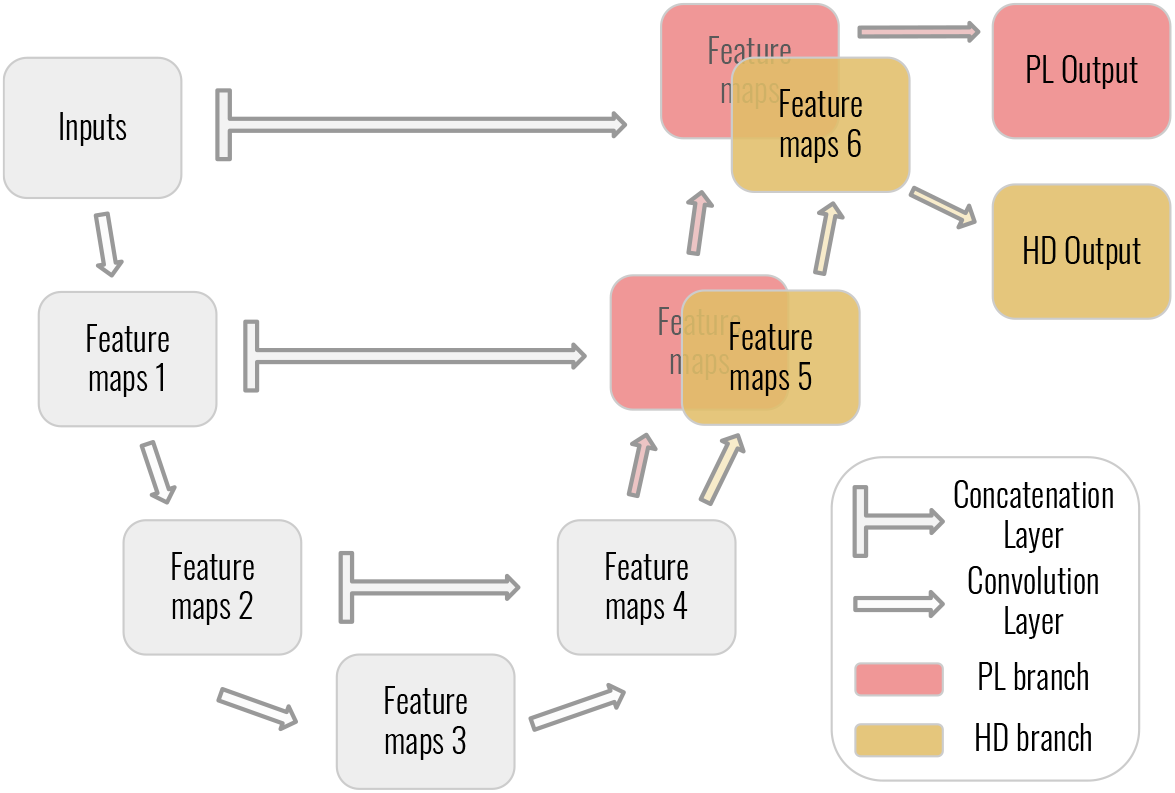
Visual representation of `InDeep`’s architecture.

We then train the resulting models with batches containing a mix of `PL` and `HD` data points. For memory limitations issues, we used the accumulated gradient trick to use batches of size greater than one. Since we encode the 3D structure into voxels, the resulting representation is sparse. Moreover, the key part of the prediction lies in the space surrounding the protein surface and especially around the position of the true ligand, while having a hard zero inside the protein of very far from its surface is less relevant. To account for these two points, we use the weighted versions of cross entropy (CE) and binary cross entropy as the loss ℒ to train our network. Voxels all receive a small weight **w**^background^ of 0.05, then the voxels closer than 6 Å from the surface receive an additional weight **w**^surface^ of 0.35 and finally the voxels corresponding to the target voxels get an additional value **w**^partner^ of 1. We found this weighting scheme to stabilize the learning and optimized our network with an Adam[47] optimizer.

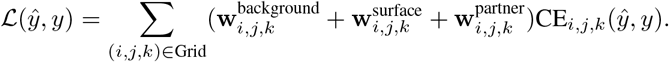

### 2.4 Post-processing

Once equipped with scalar fields prediction, we need to segment them in contiguous regions of high values. Other approaches have simply used a mean-shift algorithm [23, 25], but such algorithms can split the prediction, or discard important neighboring parts of the prediction. To address this segmenting problem, we have relied on the watershed[48] algorithm. The watershed algorithms finds all basins around local minima. We build a graph whose nodes are these basins and whose edges contain the euclidean distance as well as a normalized value of the lowest point joining two neighboring basins. Then, we merge the neighboring nodes in a greedy manner, by prioritizing the ones with the smallest edge. We stop the merging process when the merged nodes exceed a geometric distance threshold of 15 Å and 20 Å) for `PL` and `HD` respectively. Each of the resulting group of basins is denoted as a predicted pocket and scored based on the mean values of its best 150 voxels. Finally, we filter these predicted pockets to remove the smallest or least high-scoring ones, yielding the final list of predictions.

### 2.5 Hyper-parameters optimization

To choose the optimal hyperparameters, we conducted an hyper-parameter optimization (HPO). The idea is to choose these hyper-parameters (number of layers, number of neurons per layer) automatically by considering the training as a function evaluation. The function needs to output a scalar value that we want to minimize. Then, many discrete optimization algorithm such as Gaussian processes [49] can be used to find optimal parameters with regards to the minimization of this metric. We designed a custom metric to take into account several aspects of the prediction, such as the distance between a prediction and its supervision but also their overlap or the size of the prediction. This metric and optimization procedure is described in detail in the Supplemental (section C. We then ran the HPO for approximately 100 experiments, minimizing this metric on the validation set. This gave us our final predictive model.

### 2.6 Molecular Dynamics Analysis

To distinguish the conformations with higher ligandability (`PL`) or interactability (`HD`) propensity, the developed model can then be used on an ensemble of protein conformations generated by molecular dynamics (MD). To do so, each snapshot of a molecular dynamics simulation has been used as an input for `InDeep`. Therefore one can specify some residues to `InDeep` along which to monitor the ligandability or interactability, resulting in a reduced grid compared to an inference on the whole protein. Then, the post-processing is simplified because the prediction size is reduced : a spatial anchor is chosen either by the user or as the point of the grid in the solvent closest to the grid center. Finally, we simply grow a volume around this anchor following a greedy nearest neighbor policy. The average value of the voxels in this volume represents a ligandability or interactability score that can be easily tracked along the MD time steps. We chose a value of 150 Å^3^, close to the cutoff of 100 Å^3^ used in the study by Gao et al. which considers that 80% of pockets occupied by ligands are encompassed by this cutoff.

## 3 Results

### 3.1 Prediction of ligandable binding sites

#### 3.1.1 Benchmark of InDeep on conventional target binding sites

To our knowledge, there is no tool dedicated to predicting iPPI binding sites. There are however several tools that aim to predict small molecule binding sites [19, 23, 50], among which Kalasanty [25] is a reference is the field. The other major difference between our tool and Kalasanty, is that Kalasanty was trained on VolSite [51] predicted cavities in ligand locations, whereas our model has been trained on ligand position directly. In the absence of other iPPI dedicated tool for ligandability, we benchmarked `InDeep` against Kalasanty. For fairness, we compare the ability of our tool to predict VolSite predicted cavities as well as the ligand pose with Kalasanty.

We use the same dataset as the authors of Kalasanty did for validation: a distinct dataset, made by Chen *et al* [52]. The original test set is composed of 111 protein-ligand holo structures and 104 corresponding apo structures. We have filtered out a few systems that were too similar to our training set according to the TM-score metric [53], as shown in Figure 2 and end up with 187 and 196 pockets evaluation for apo and holo structures respectively. Ligands coordinates were extracted from the holo structures of the Chen benchmark and VolSite [51] was used to describe cavities for each ligand. Following Kalasanty, the number of retained predicted pockets is the number of small molecules present in the deposited PDB system.

**Figure 2:**
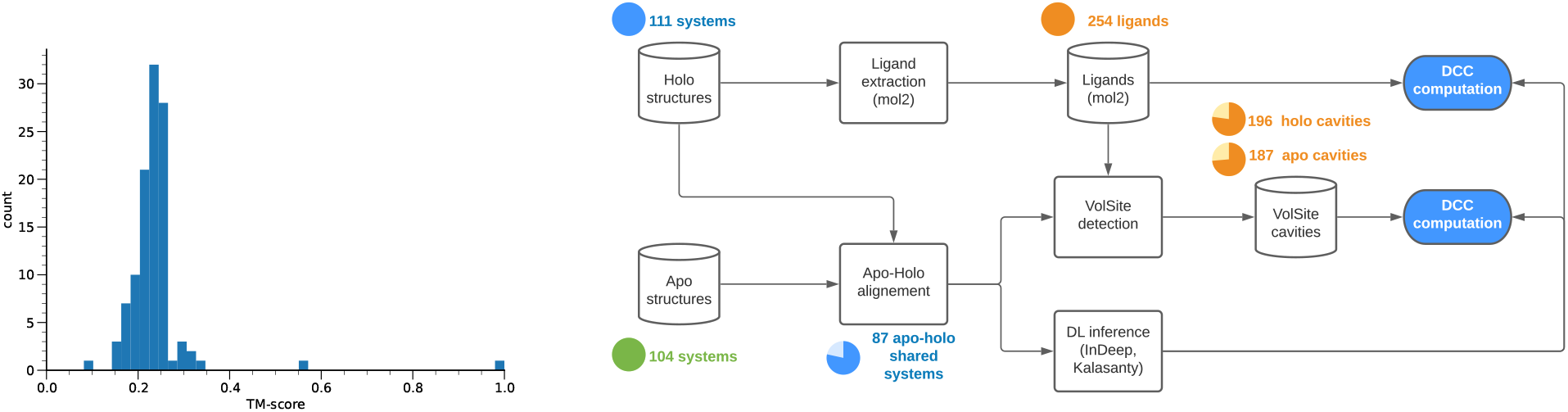
*Left* : TM-score between the Chen benchmark and the `InDeep` training set. *Right* : Overall process of the metric calculations for `PL`. Figures correspond to the Chen benchmark.

We then used the metrics used by Kalasanty on their dataset: DCC and DVO. The DCC metric computes the distance in Angstrom between the center of mass of the prediction and the one of the ground truth. We denote a prediction as successful when its DCC is below a distance threshold, and we plot the success rate of the different methods. We compare predictions of `InDeep` compared to the ones of Kalasanty, for the bound conformation (holo) as well as the unbound one (apo). The DCC values were computed in 3 conditions: between the computational predictions and (i) the VolSite cavities, (ii) the ligand positions and (iii) the ligand positions that have a VolSite cavity associated with them. This procedure is illustrated in Figure 2 and the results are presented in Figure 3a A, B and C respectively. The DVO metric is only computed on successful prediction at 6 Å and consists in the volume of the overlap over the volume of the union (Figure 3b).

**Figure 3.**
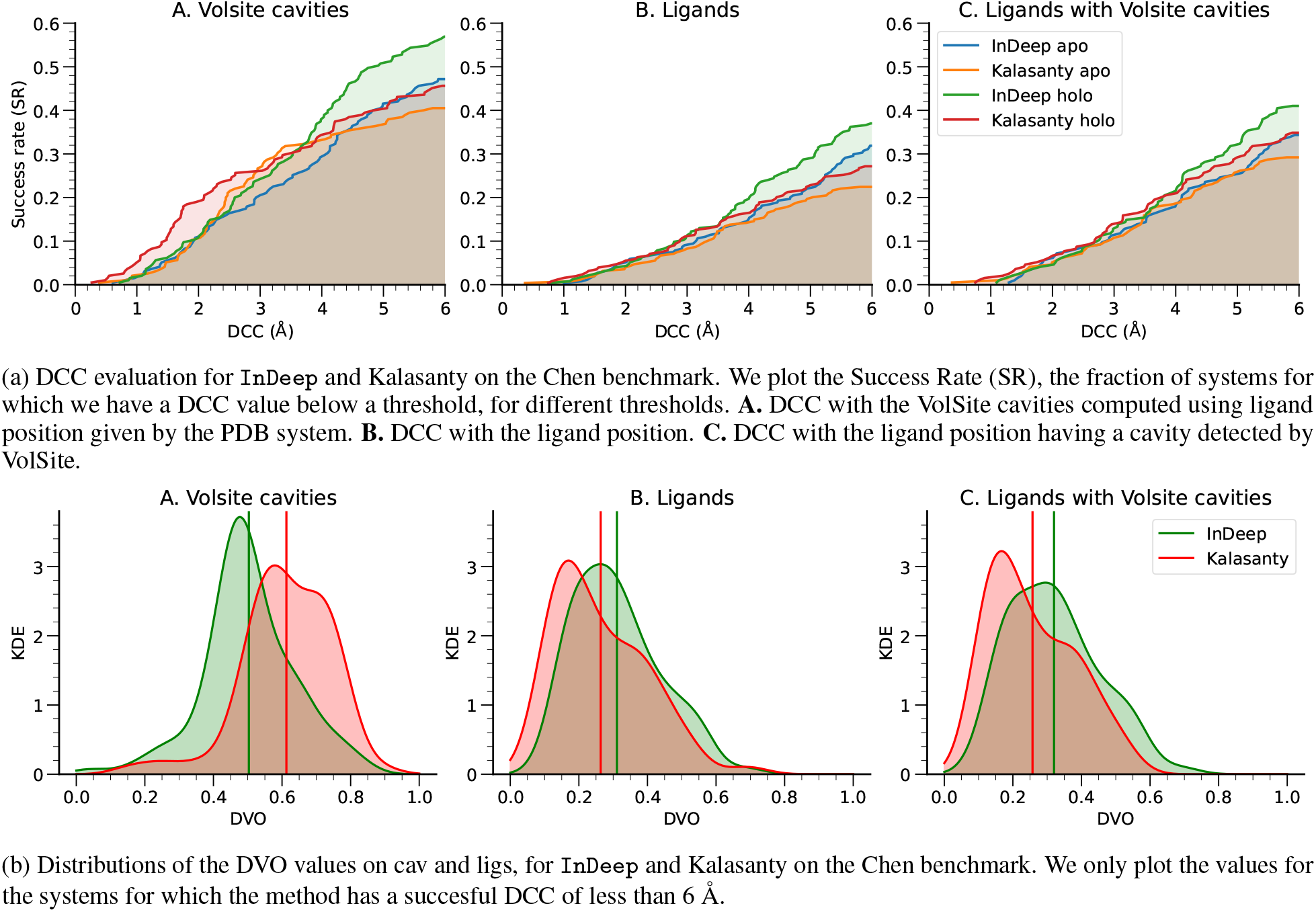

First of all, we reproduce the results claimed by Kalasanty. In this benchmark dataset, `InDeep` outperforms Kalasanty on all settings except in the very low DCC range for cavities. At a 6 Å threshold, we have an average relative performance boost of 27%. We see that this difference is most important in the ligand setting that resembles our training procedure, but that the performance remains stable on the cavities setting. We also see that this boost of performance is observed across holo and apo predictions. We detect approximately 40% of the binding sites in the best ranked predictions. We then compute the DVO values for the different methods among the successful predictions at 6 Å. We obtain comparable values of DVO overall. Because Volsite cavities tend to be deeper into the pocket, Kalasanty predictions tend to be at the bottom of binding sites. This explains why the average DVO value for cavities is better for Kalasanty and worse for `InDeep` (0.612 vs 0.503), and vice-versa for ligands (0.257 vs 0.320). Moreover, one should note that the total populations are not the same because the DVO is computed only on the ’successful predictions’ with low DCC values. Therefore, the better performance in terms of DCC can hinder the DVO distribution performance. Overall, `InDeep` yields state of the art performance on the binding site prediction for general ligands, even in the unfavorable setting of predicting cavities.

#### 3.1.2 Benchmark of InDeep on iPPI binding sites

We have then repeated the same data extraction pipeline as used for the Chen data set and described in Figure 2 on our test test. We have applied the same TM-score filtering between the train set of Kalasanty and our test set to avoid data leakage. This represents a more suitable application InDeep, as it was designed to identify PPI specific binding pockets.

We also note that this procedure results in few (81) systems, because of the large training set of Kalasanty. We present the DVO and DCC result we obtain in Figure 4.

**Figure 4:**
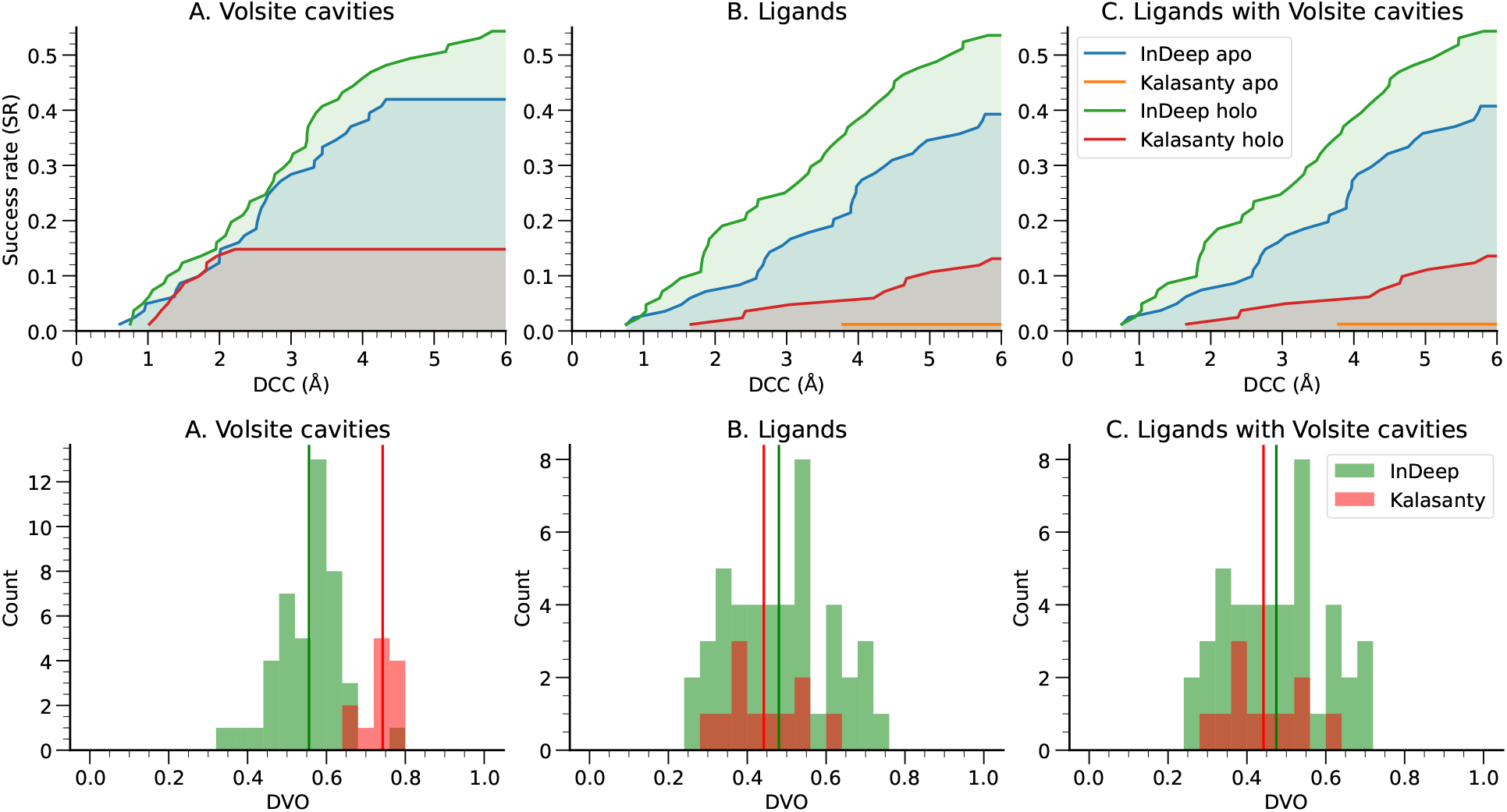
Performance on the test set filtered by TM-score. The plots are produced following the same procedure as the ones above on a the new data set.

Despite the limited size of this filtered data set, we see that `InDeep` clearly outperforms Kalasanty. The SR is increased 5-fold and the DVO values remain reasonable. This shows that `InDeep` is not only a good predictor for conventional target binding sites, but that it is much more efficient than existing methods for iPPI binding site detection.

### 3.2 Predicting and profiling epitope binding sites

#### 3.2.1 Benchmark of InDeep on PPI data sets

As for `PL`, several tools exist that predict which region of a protein interact with another. Once again we choose to compare against the state of the art and reproduce the results of PInet. We use two benchmarking data sets they propose : DBD5[54] and EpiPred[55]. DBD5 is a protein docking benchmark that offers several pairs of structures of interacting proteins. EpiPred is a dataset centered around interactions between antigens and antibodies.

We then had to slightly adapt our validation pipeline. Indeed, we have found no study trying to predict the actual location of the partner in the vicinity of the protein surface. The tools we compare against always project the predicted interactability onto the surface residues of the protein or the sequence. Therefore to compare against other tools, we needed to implement a sequence supervision and to project our 3D prediction onto the sequence. To do so, we use a convolution with Gaussian kernel between our 3D prediction and the coordinates of the atoms of the protein. We optimized the parameters of the projection, such as the kernel radius, using a grid search on the DBD5 train set. We then use those parameters on the DBD5 test set as well as the EpiPred data set. For fairness, we used the same method to annotate the data as in PInet and we have rerun their method. Finally, we have used their tool in the partner-specific setting (giving the partner as input) and in a blind setting (PInet blind) that is closer to our use case. The results are presented in Table 1.

**Table 1:** Area under the Precision-Recall curve for binary prediction of interaction on the sequence, on the DBD5 and EpiPred data sets, for `InDeep` and PInet.

| Dataset/Method | PInet | PInet blind | InDeep |
| --- | --- | --- | --- |
| DBD5 | 0.707 | 0.647 | 0.3154 |
| EpiPred | 0.235 | 0.217 | 0.2319 |

We see that on the DBD5 test set, `InDeep` is outperformed by PInet. However, PInet was trained on the same data set, and we turn to EpiPred, that was not used for training by either methods. On this data set, our result falls close to the performance of the partner-specific setting. We achieve a state of the art performance in our blind setting. Moreover, it should be noted that `InDeep` performance suffers from the extra step of sequence projection. Overall this shows that our tool is able to accurately predict the interaction sites of a protein.

#### 3.2.2 Localization of interacting partner

We complement this comparison to other tools with a validation of `InDeep` with metrics closer to the `PL` validation. We note that to our knowledge, no tool exist that output 3D prediction of the volume occupied by a putative protein partner. However, this prediction is of a great use to assess if the 3D prediction for a small molecule binding would collide into its corresponding protein partner, opening new doors for therapeutic design of iPPIs. Since for PPIs, the number of observed partners is just one, we have computed DCC and DVO values for one, three and all predicted binding sites. Because the interfaces are bigger, we also present our results up to a distance threshold of 10A. The results are presented on Figure 5.

**Figure 5:**
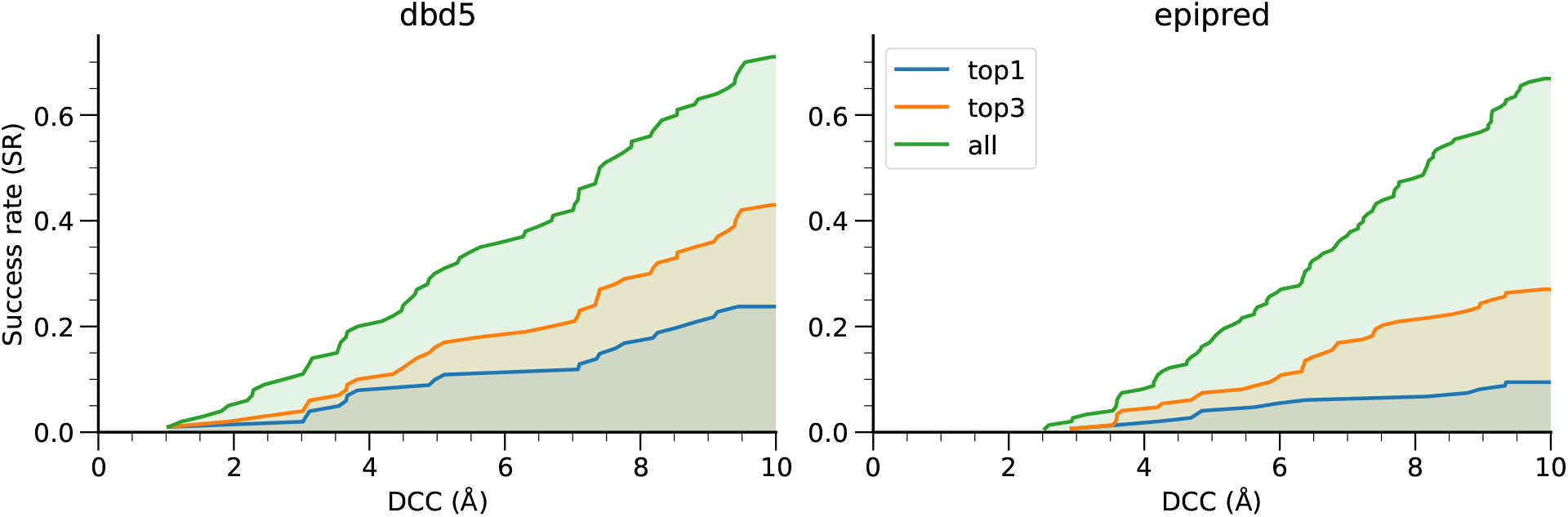
DCC values for `InDeep` on DBD5 (*Left*) and EpiPred (*Right*) data sets, when considering the best, top-3 or all pockets predicted.

At the 10 Å threshold, we have success rates of 24% and 11% with only the first prediction, and of 72% and 69% using all of them for DBD5 and EpiPred respectively. This means that our method finds the correct binding site in of 70% of the cases, but that a significant amount of times, the correct predicted volume is not ranked as the first one. This can be partially imputed to the fact that a given protein can have several partners, so the first prediction might actually be a correct one that does not correspond to the partner at hand. This is the motivation for partner-specific prediction tools, such as PInet. However, `InDeep`’s performance on the top-3 falls in between the performance of the top-1 and keeping all pockets. This indicates that the correct binding sites is often among the best scored position, proving once again the relevance of the tool for epitope binding site prediction.

#### 3.2.3 Epitope binding site prediction: atom typed channel validation

We now turn to the channel evaluation. We have used five channels to encode our protein environment and our prediction : α-carbon (’CA’), donor and receptor of h-bonds (’HAD’), positively and negatively charged atoms (’POS’ and ’NEG’) and hydrophobic/aromatic atoms (’COB’).

This means that beyond prediction of the presence or absence of a protein partner, the model also predicts which protein atom type should be at a given voxel. This is close to the idea developed by LigVoxel[50] but for the interactability model. However, finding a quantitative metric to describe the quality of these channels is not easy. Indeed, the target now contains several little volumes (each atom type environment) that can be split across the protein partner interface. Therefore, we can not use the DCC metric easily, because the centers of mass of split volumes do not represent our objects accurately. Moreover, we cannot easily interpret the DVO values, as previous experiments only plotted the DVO for successful DCCs, which we do not have anymore. This is even more true if we consider the large size of the interface, which explains why we cannot use the same validation procedure as LigVoxel[50].

We have turned to a more direct method for assessing the performance. At each voxel of our prediction, we have a distribution of probability for each channel. We can aggregate these voxel distributions for all voxels around an atom of the ligand to obtain a mean distribution of channels probabilities. We also compute an atom-type specific distribution by aggregating only the voxels around atoms of each specific channel. We plot a heatmap representing the Z-scores of the observed channels distributions compared to the overall ones. We expect to see enhanced values on the diagonal and decreased ones off the diagonal. The results are presented in Figure 6.

**Figure 6:**
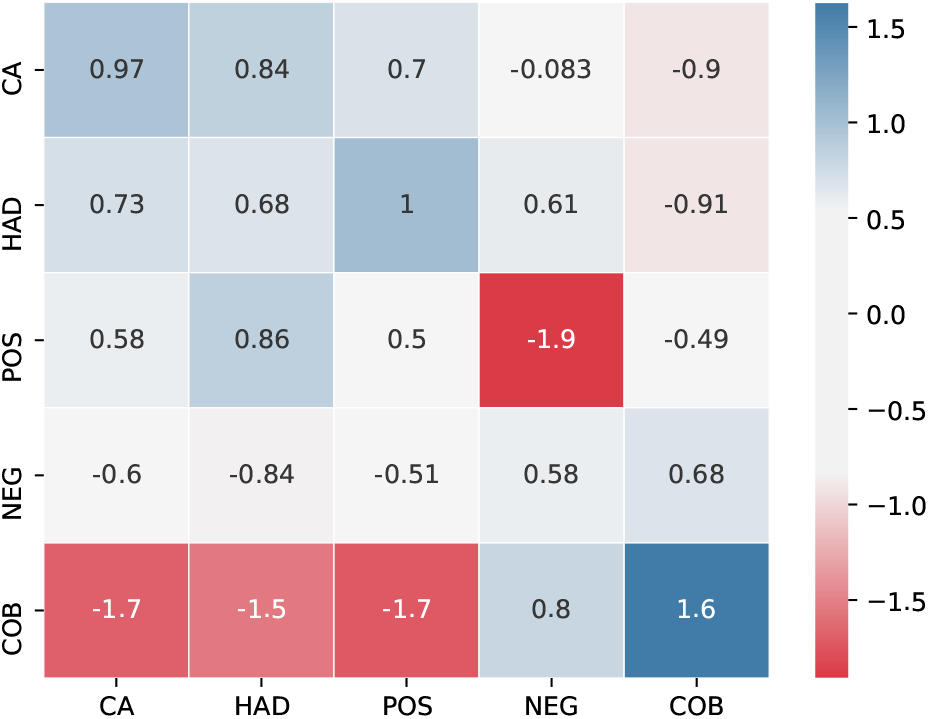
Z-scores of the predicted distributions of the probabilities for each channel (one distribution per line). Each line correspond to a distribution around atoms of the partner that have a certain channel annotation.

We see that the hydrophobic channel (COB) performs well at localizing hydrophobic patches of protein partners (Z-Score = 1.6). It is an important result as transient protein-protein interactions, that represents most of the known PPI targets, are often mediated by hydrophobic patches at the interface whose seclusion from the solvent upon binding helps to regulate protein association. This COB channel can therefore be used as a way to suggest point mutations at the interface when dealing with hydrophobic interaction, in the context of epitope mimicking or peptide design. The backbone channel (CA) has a more modest performance than COB although it displays a partial enrichment. In this case, the perspective of depicting the backbone of a putative partner for a given interactability patch is also very pertinent. Indeed, for example the spatial arrangements of Cα within a α-helix are fitting very nicely within a cylinder that can be clearly identified within some PPIs mediated by such secondary structure at the interface (see case study about Bcl-2). Nevertheless, the other channels are clearly non specific and shall be the subject of improvements in the future.

### 3.3 Case study: Bcl-2 as therapeutic target

The B-cell lymphoma-2 (Bcl-2) protein is the eponymous protein of the Bcl-2 family which is central to the regulation of apoptosis and vital for proper tissue development and cellular homeostasis [56]. Upon interaction with pro-apoptotic BH3 domains containing proteins, Bcl-2 inhibits cell death. In the last two decades, small molecules that disrupt this interaction by binding, to Bcl-2 and other anti-apoptotic proteins of this family, have been successfully designed and clinically approved to induce apoptosis of cancer cells [57]. Pro-apoptotic partners of Bcl-2 possess a 20 Å-long α-helical-containing BH3 domains that interact at Bcl-2 surface through an extended hydrophobic groove. A recent review underpinned that the successful development of drugs, such as Venetoclax, against this target was mainly due to the fact they manage to mimic two of the hotspot residues within the BH3 domain binding this groove [4].

This protein family is therefore an excellent case study to evaluate the pertinence of using a tool like `InDeep`. The fact that this tool can predict ligandable/druggable binding pockets, interactability patches including hydrophobic and backbone atom-typed channels, and also monitor such predictions along molecular dynamics trajectories allows a retrospective analysis of feasibility of designing ligands binding to the BH3 groove of Bcl-2. It is worth noting that the different `InDeep` predictions below have been made exclusively on the sole structure of Bcl-2 without any consideration for Bax or known co-crystallized ligands.

We can first use `InDeep` to predict interactability patches at the surface of Bcl-2. As can be seen on Figure 7, `InDeep` correctly predicts (1st ranked patch), within the BH3 groove, the location of the interactability patch with the α-helix of its protein partner Bax (Left panel). Inspecting more specifically the Cα- and hydrophobic-atom-typed channels within the interactability patch (Right panel), one can observe respectively 1) a faithful depiction of the α-helix shape of the Bax epitope binding the BH3 groove (as a cylinder shape), and 2) a proper localization of the hydrophobic hotspots known for this system [4]. It is important to note that all the `InDeep` predictions were performed only on Bcl-2, without any knowledge of protein partners or ligands binding information.

**Figure 7:**
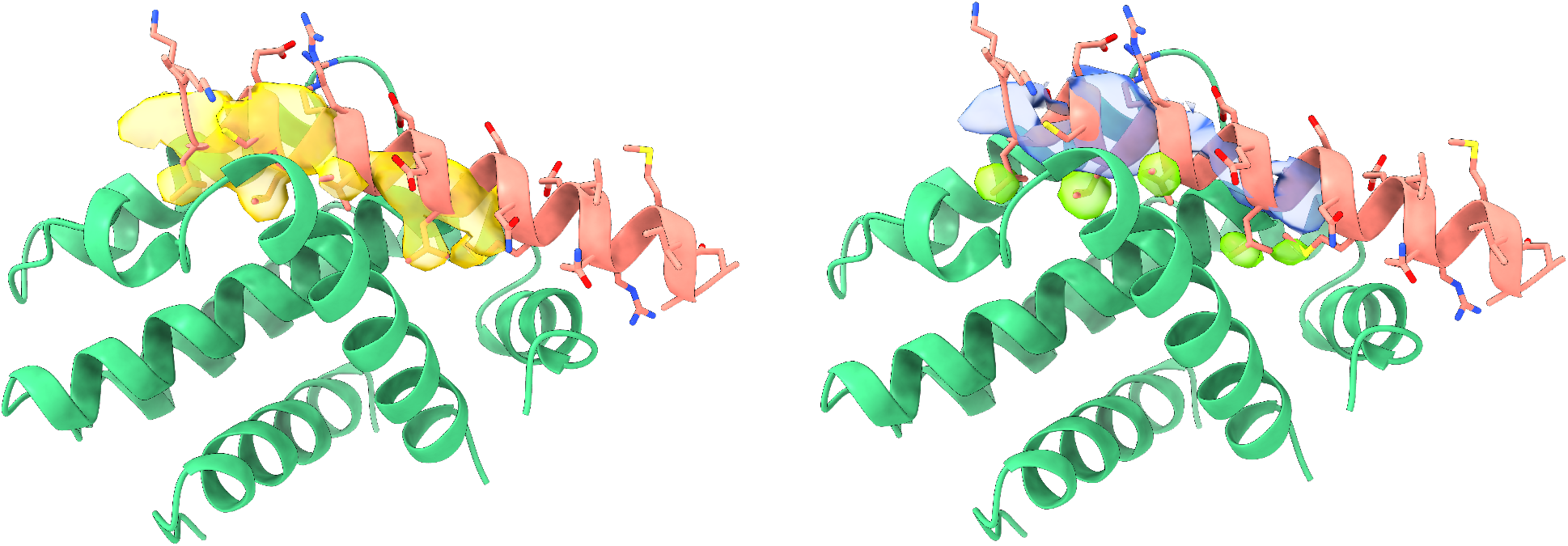
*Left* : `InDeep` interactability patch prediction on Bcl-2 (pdb 2xa0). *Right* : The hydrophobic-atom typed channel (green surface), predicted on Bcl-2, (green cartoon) correctly matches with the known hydrophobic hotspots of the protein partner Bax (orange cartoon). Likewise, the Cα-atom typed channel (blue surface) correctly predicts the α-helix shape of the Bax backbone (depicted a shape close to a cylinder).

If we now use the ligandability prediction of `InDeep` on the same structure of Bcl-2 as co-crystallized with Bax (pdb: 2xa0), one can see on Figure 8 that `InDeep` correctly highlights the known aforementioned hotspots as the most ligandable regions of the BH3 groove of Bcl-2. The same regions that were indeed targeted by Venetoclax analogs (ex with pdb: 4lvt).

**Figure 8:**
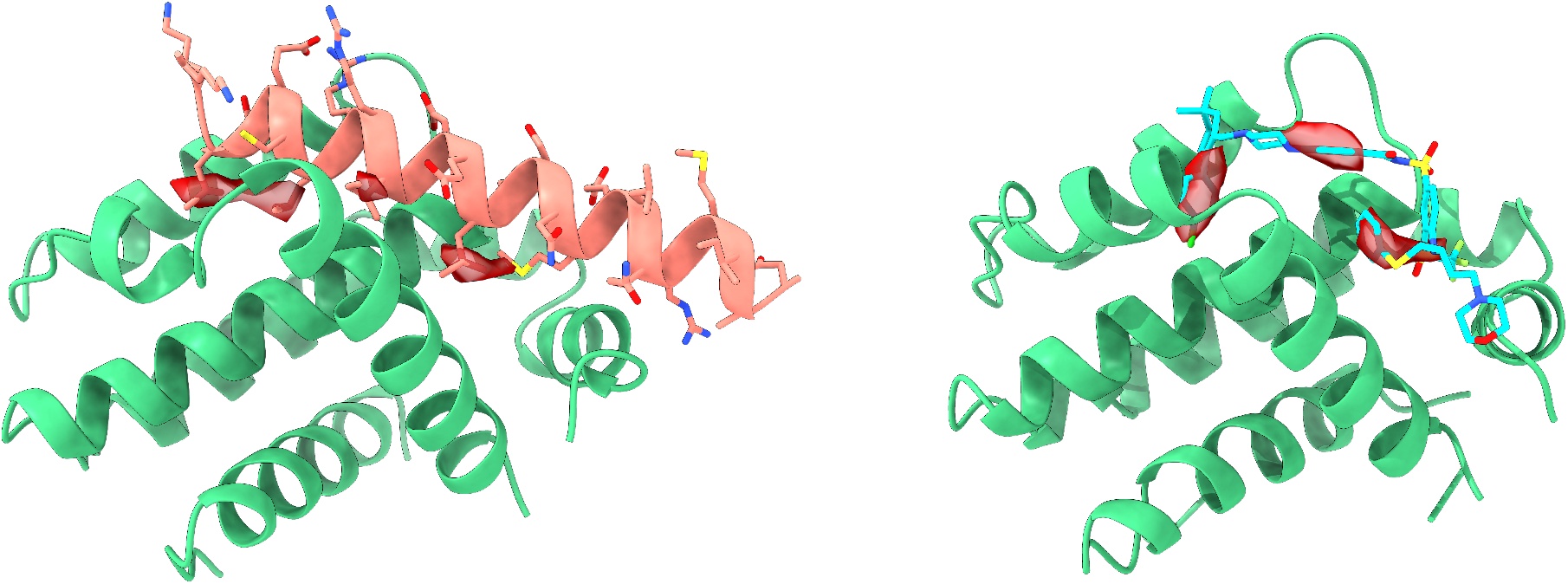
*Left:* `InDeep` Ligandability prediction on Bcl-2 (pdb 2xa0). The red surface patches of `InDeep` ligandability are localized around the known hot spots of the Bcl-2/Bax interaction. *Right:* Ligandability prediction (red surface) performed on Bcl-2 (pdb 4lvt) surface. The red surface patches of `InDeep` ligandability are localized around the known hot spots of the Bcl-2/Bax interaction that are mimicked by some of the ligand atoms.

NMR and X-ray crystallography have highlighted important backbone rearrangements within Bcl-2 upon peptide and small molecule binding to the BH3 groove [58]. Early virtual screening approaches failed to identify validated small molecules inhibitors as they did not consider protein flexibility [59]. It is therefore essential to properly sample the flexibility of the system and be able to monitor both ligandable and interactability patches on various Bcl-2 structure conformations. To do so, a 1 µs-long molecular dynamics simulation (see MD parameters in Supplemental section E), starting from the apo form of Bcl-2 (pdb 1gjh), was run to monitor the ligandability and the interactability of the BH3 groove known to bind Venetoclax and other chemical analogs. The DL prediction has been focused on this region of Bcl-2, as the goal of this approach is to detect holo-like conformations i.e favorable conformations for ligand binding and not to detect the binding region on the whole protein surface. Figure 9 (Left panel) shows the values of the `InDeep` ligandability score of each frame and the minimal binding site RMSD value with respect to several ligand-bound `PL` structures (see binding site definition and the list of `PL` structures in F. It can be noted that holo-like conformations (with low RMSD against a `PL`) tend to have high ligandability scores. For example, at around 500 ns of simulation, a decrease of the RMSD, and therefore an increase of holo-likeness, is clearly detected by `InDeep` with an increase in the ligandability score. We thus confirmed, on a PPI target using `InDeep`, what other tools could observe on conventional targets [36, 37]. Likewise, the interactability was monitored along the same simulation and the RMSD of the binding site residues was compared to a holo `HD` structure of Bcl-2 bound to Bax (pdb 2xa0) Figure 9 (Right panel). Similarly to the ligandability model, holo-like conformations are better scored by `InDeep` interactability model than conformations far from the `HD` conformations.

**Figure 9:**
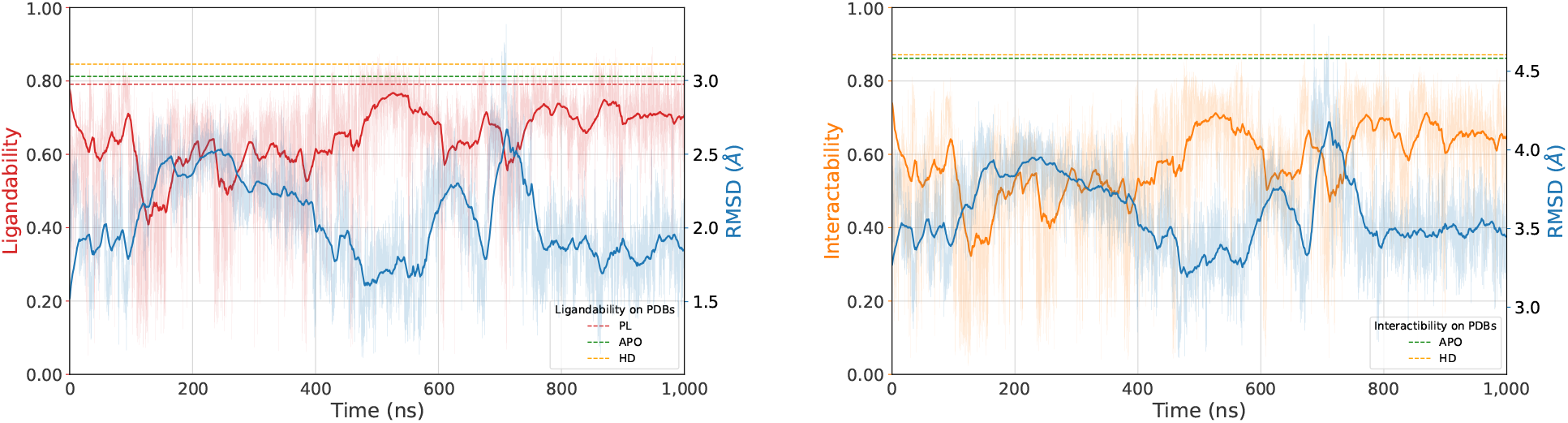
`InDeep` predictions (red and orange) along molecular dynamics trajectory of Bcl-2. Moving average (exponential) on 500 frames are represented with solid lines. *Left* : `InDeep` ligandability score evolution (red) compared to the minimal binding site RMSD (blue) with respect to 16 `PL` structures. Red horizontal dashed line represents the average ligandability of the 16 references `PL` structures (0.79*±* 0.04), green horizontal dashed line represents the ligandability of the apo starting point of the MD and orange dashed line represents the ligandability of the `HD` structure. *Right* : `InDeep` interactability score evolution (orange) compared to the RMSD (blue) with respect to the reference `HD` structure. Orange horizontal dashed line represents the interactability of the reference `HD` structure and green horizontal dashed line represents the interactability of the apo starting point of the MD.

These predictions collectively show that a proper usage of `InDeep` early in the drug discovery initiative against Bcl-2/Bax would have highlighted the hydrophobic hotspots residues within the BH3 groove of Bcl-2 and the most ligandable binding regions of these spots to assist the design of selective BH3 mimetics. Moreover, `InDeep` would have been efficient at profiling holo-like conformations prior to virtual screening campaigns even when starting from apo structures, the absence of Bax bound to Bcl-2, and within a molecular dynamics trajectory.

## 4 Discussion

We have introduced `InDeep`, a unified prediction tool for structure-based drug design targeting protein interfaces. We show that this tool is competitive in detecting the residues that interact with a protein partner. We go beyond this sequence prediction by predicting the 3D shape of the output with atom-typed channels signal that helps understanding how such protein interaction can take places. 70% of the observed binding site are present in one of our prediction and 35% in the top three ones. Moreover, `InDeep` outperforms the state of the art of binding pockets detection for conventional targets and we showed that our tool is even more efficient to detect iPPI binding sites. Combining those two predictions for a newly solved structure, one can investigate binding sites for ligands that would potentially disrupt a given PPI. Given such a detected binding site, our tool also enables tracking ligandability or interactability scores along a molecular dynamics trajectory, which opens the door for a refined ligandability assessment as well as conformation selection for virtual screening, or unravel epitope binding-prone conformations. Finally, we illustrate these functionalities in a retrospective drug discovery use case on Bcl-2. `InDeep` is integrated in a PyMol [38] plugin for easy visualization of the predictions (see Section D).

Despite several promising results, the therapeutic use of iPPIs remains a minority. We hope this dedicated tool can help enhance their use as well as spark a development of other methods following this line of work. `InDeep` uses 3D-FCNs because the grid-like prediction enables actual localization and profiling of protein partners or future drugs. However this method, as many others, is sensitive to rotation, a possible extension would be to use equivariant networks [60, 61, 62] to bake the rotation invariance in the network.

Future developments also include actual drug discovery project use of the tool as well as the implementation of a web-server for easier access to the predictions.

## 5 Acknowledgements

We thank IBM for their sponsorship for the optimization of the hyperparamaters. In particular, we thank Jean Armand Broyelle, Maxime Deloche, and Xavier Vasquez for their expert support. We are grateful to the Institut Pasteur and the CNRS for their continued support for our research. V. M. is recipient of a doctoral fellowship from the the INCEPTION project [PIA/ANR-16-CONV-0005] and benefits from support from the CRI through “Ecole Doctorale FIRE – Programme Bettencourt”. The authors also thank Arnaud Blondel for feedback and discussions. They declare no conflict of interest.

## A Data splitting

To avoid data leakage, we split our protein structures data based on CATH[40] folds. We have excluded the following CATH classes from the train split : 1-25-10-10, 1-10-530-10, 1-10-245-10, 1-20-120-560, 1-50-10-20, 2-30-29-30, 2-30-42-10, 2-130-10-10, 2-60-120-620, 2-30-30-50, 3-40-390-10, 3-30-40-10, 3-90-70-10, 3-10-110-10, 3-90-540-10.

## B Model

The model consist in blocks that make two 3D convolution, batch normalizations and ELU activations. The first convolution goes from in_channels to `(in_channels+out_channels)/2` and the second goes to `out_channels`. Following the classical structure of a UNet model, at each level, we use one such block to decrease the spatial extension of the block by a factor two, while doubling the channels. We start with eight channels and use a depth of four. We use a twist on the classical UNet, because we branch our network into a `PL` and `HD` one at the last deconvolution block of the decoder. Then each branch get convoluted one last time and is activated by a sigmoid or a softmax.

## C HPO

In this section, we describe the procedure used to tune hyperparameters of our method. Hyper-Parameter Optimization (HPO) consists in finding parameters that optimize a given scalar metric.

### C.1 Metric

The metric we chose to optimize need to account for :

- The accurate position of at least one predicted volume
- A good ranking of our correct predicted volume
- A good shape correspondence between the predicted volume and the actual partner

We first compute a distance metric called dist_overlap that is the mean of the distance between the center of mass of the predicted volume and the partner and the overlap between the highest scoring voxels of the predicted volume and the partner. We then pick the predicted volume that yields the best dist_overlap value and compute two additional metrics to encourage giving it a high rank. The first one is simply the exponential of its rank over its surface (to normalize the rank by the expected number of volumes). The second one, deemed as rank_overdist is an expectation-like formula as follows : 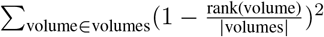 dist_overlap(volume). We add a DVO score for the shape complementarity. Finally, we add a penalty for having too many predicted volumes by subtracting to the metric value the number of predicted volumes divided by the surface.

All of these contributions are then summed and we empirically checked on the validation set that the satisfactory looking predictions had a high metric value while failed ones had a low value. We can then proceed to the automatic optimization of the hyperparameters with regards to this scalar metric.

### C.2 Hyper-optimization

We have tweaked the type of model (U-Net vs forward), the type of blocks used by the UNet, the amount of neurons at the beginning at the network and the depth of the network. Moreover, we have also tuned some post-processing parameters, namely the maximum euclidean distance between two watershed basins and the threshold of the merging value between two neighboring basins.

The convergence of this procedure is plotted in Figure 10.

**Figure 10:**
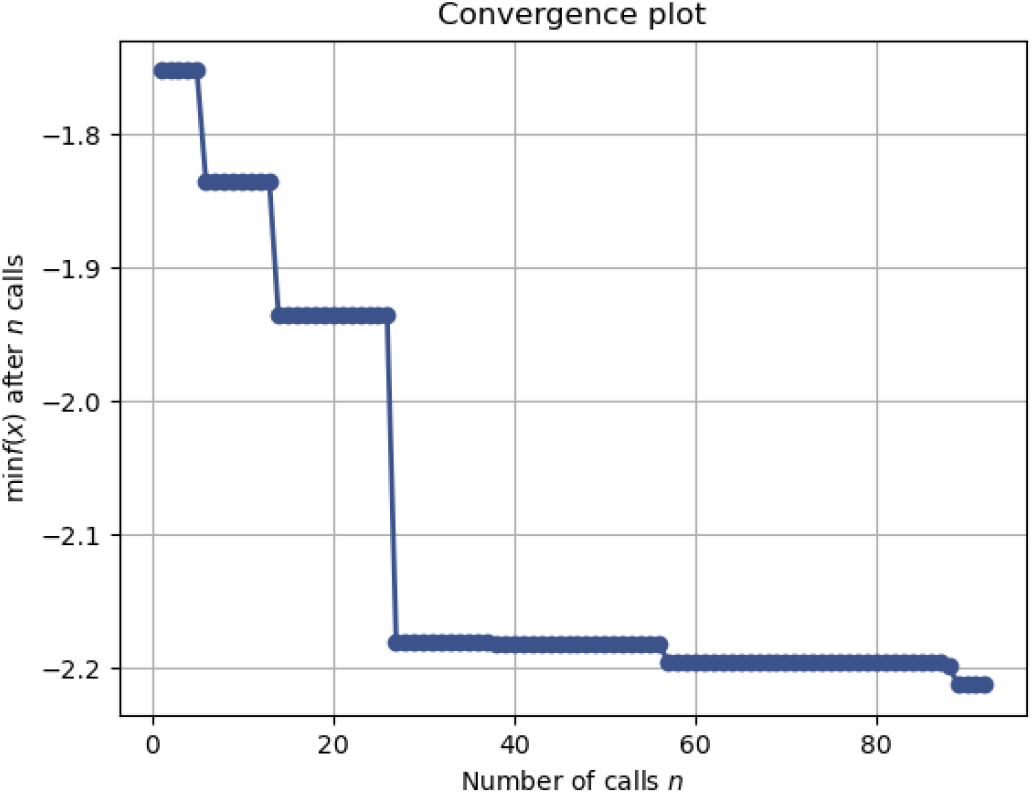
Successive values of the HPO metric as a function of the HPO epochs

## D Pymol plugin

The tool can be added as a PyMol plugin for an interactive visualisation of the results. A screenshot is presented in Figure 11.

**Figure 11:**
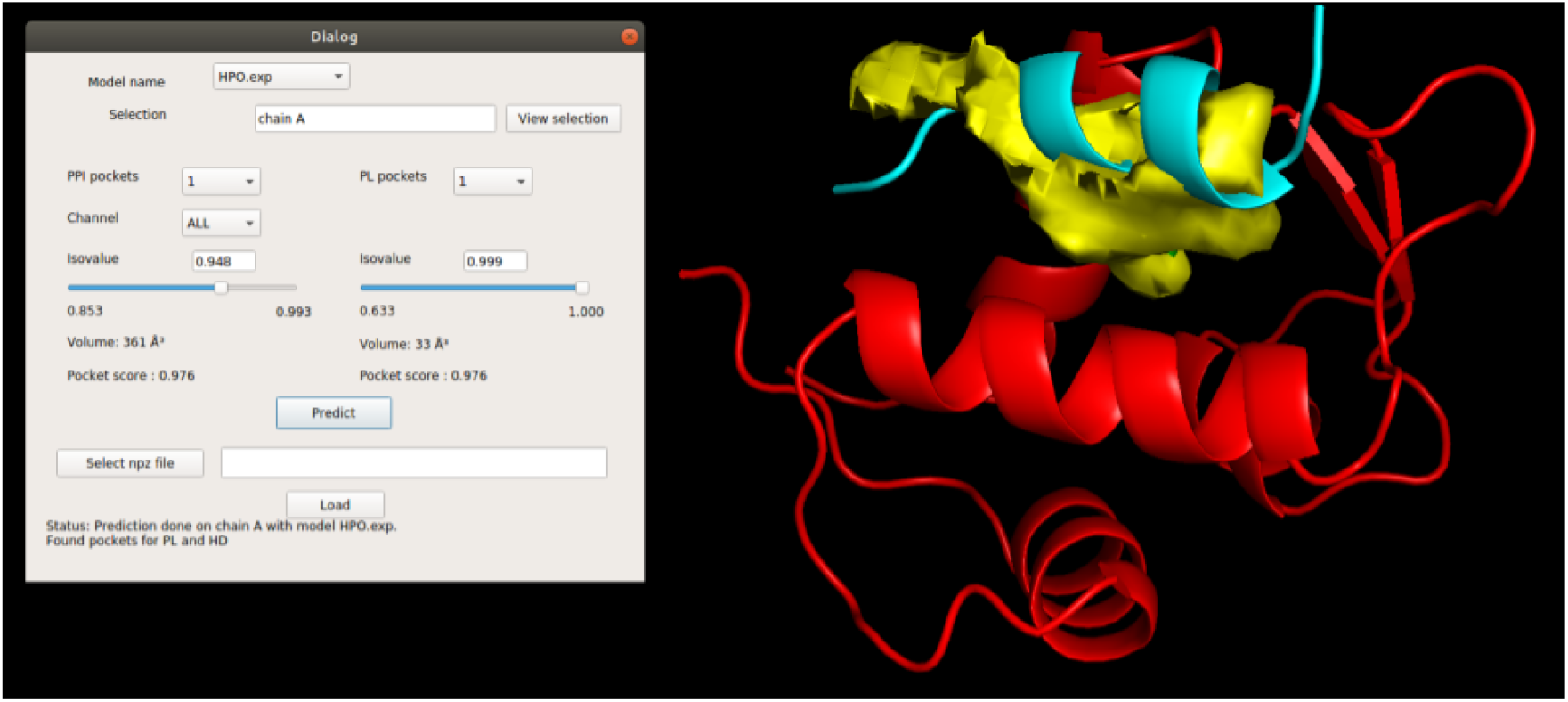
The results can be visualized through a PyMol plugin, that lets the user see the different pockets and their corresponding volumes at different probability levels

## E Molecular Dynamics setup

The MD input files were prepared from the unbound form of Bcl-2 (pdb 1gjh) with CHARMM-GUI solution builder server [63] and CHARMM36m force field [64]. The protein was placed in a cubic unit cell with a minimum distance of 10 Å to the box edge. The system was solvated with an explicit water model (TIP3P) and KCl ions were added at 0.15 M concentration. Steepest descent minimization was performed in 5000 steps. Subsequently, a 125 ps equilibration in the NVT ensemble was performed at 303.15 K. Finally, production was run for 1µs in the NPT ensemble. GROMACS [65] was used for the simulations.

## F Binding site definition for RMSD calculations and InDeep predictions during MD

In order to obtain the list of residues defining the Bcl-2 orthosteric binding site we have retrieved from the dataset 16 `PL` structures of Bcl-2: 1ysw, 2o21, 2o22, 2o2f, 2w3l, 4aq3, 4ieh, 4lvt, 4lxd, 4man, 6gl8, 6o0k, 6o0l, 6o0m, 6o0o and 6o0p. Then, for each structure the solvent accessible area by residue was measured with naccess [66] in presence and absence of the orthosteric ligand. Residues having lower solvent accessible area when the ligand is present are considered aspart of the binding site. As a result the following residues were used to define the binding site: 96, 99, 100, 103, 104, 107, 108, 111, 112, 115, 133, 136, 137, 144, 145, 146, 148, 149, 152, 153, 156, 198, 202 and 143. Heavy atoms of these residues have been selected to measured the RMSD along the MD compared to the 16 `PL` structures and to the `HD` structure (pdb 2xa0). The reuslting RMSD curves are shown in Figure 9.

## Footnotes

3 Code available at : https://gitlab.pasteur.fr/InDeep/InDeep

4 Predictions of InDeep available at : https://ippidb.pasteur.fr/targetcentric/

5 Target centric mode of iPPI-DB available at : https://ippidb.pasteur.fr/targetcentric/

## Notes

### Competing Interest Statement

The authors have declared no competing interest.

